# More Highly Myelinated White Matter Tracts are Associated with Faster Processing Speed in Healthy Adults

**DOI:** 10.1101/152546

**Authors:** Sidhant Chopra, Marnie Shaw, Thomas Shaw, Perminder S Sachdev, Kaarin J Anstey, Nicolas Cherbuin

## Abstract

The objective of this study was to investigate whether the myelin content of white matter tracts is predictive of cogni–tive processing speed and whether such associations are modulated by age. Associations between myelin content and processing speed was assessed in 570 community-living individuals (277 middle-age, 293 older-age). Myelin content was measured using the mean T1w/T2w magnetic resonance ratio, in six white matter tracts (anterior corona radiata, superior corona radiata, pontine crossing tract, anterior limb of the internal capsule, genu of the corpus callosum, and splenium of the corpus callosum). Processing speed was estimated by extracting a principal component from 5 sep–arate tests of processing speed. It was found that myelin content of the bilateral anterior limb of the internal capsule and left splenium of the corpus callosum were significant predictors of processing speed, even after controlling for socio-demographic, health and genetic variables and correcting for multiple comparisons. A 1 SD increase in the myelin content of the anterior limb of the internal capsule was associated with 2.53% increase in processing speed and within the left splenium of the corpus callosum with a 2.20% increase in processing speed. In addition, significant differences in myelin content between middle-age and older participants were found in all six white matter tracts. The present results indicate that myelin content, estimated in vivo using a neuroimaging approach in healthy older adults is sufficiently precise to predict variability in processing speed in behavioural measures.

## Introduction

The highly-myelinated nature of the human brain and the vulnerability of myelin to degeneration, may contribute to our species unique susceptibility to age-related neurocognitive disorders. The cognitive domain most associated with myelination is processing speed (PS)(Lu et al., 2011; Lu et al., 2013) a sensitive indicator of overall cognitive decline (Finkel et al., 2007; Cherbuin et al., 2010). PS can be conceptualised as the rate at which cognitive operations are executed, such as planning and initiation of intended motion and is often tested in conjunction with psychomotor speed, which accounts for the speed of the motion itself (Cepeda et al., 2013). Recentcent longitudinal studies have demonstrated an inverse U-shaped lifespan trajectory of myelin content with a peak at around 3040 years (Bartzokis et al., 2012). A simi–lar trajectory has been observed in cognitive PS scores across the lifespan (Cerella and Hale, 1994; Bartzokis et al., 2010). Further, a decline in PS is the primary cogni–tive deficit underlying the rapid cognitive decline seen in demyelinating diseases such multiple sclerosis (Demaree et al., 1999). In addition, myelin loss and PS decline have also been shown to share multiple risk-factors including APO*ϵ4 genotype (Bartzokis et al., 2009), and lifestyle factors (Anstey et al., 2009; Ramagopalan et al., 2010).

We are not aware of any study having directly exam–ined the relationship between myelin content and PS in non-clinical populations using a measure specifically de–veloped for this purpose. This is likely due to the dif–ficulty in measuring myelin levels in vivo. Histological myelin measurement is the gold standard, but it can only be performed post-mortem and is therefore not suitable to investigate this question in humans. A few studies have made important contributions in this area by using indi–rect measures such as fractional anisotropy and other dif–fusion measures. However, such measures are unspecific as they index the movement of water molecules which are affected, apart from myelin, by neuronal and glial density and size (Winston, 2012), as well as pathological states such as amyloid beta deposition (Racine et al., 2014).

Recently, a new measure, the ratio between an individu–als structural T1-weighted (T1-w) and T2-weighted (T2-w) image (T1w/T2w), has been proposed as a practical and sensitive measure for in vivo myelin content estima–tion (Glasser and Van Essen, 2011; Ganzetti et al., 2014). While not a pure measure of underlying tissue, the inten–sity of a voxel in a T1-w image is thought to be propor–tional to myelin content, whereas the T2-w image is in–versely proportional. As such, a ratio between the images produces an image that has enhanced contrast sensitivity to myelin and that attenuates most scanner biases. Re–cent attempts to validate the T1w/T2w have demonstrated a concordance with immunostaining for brain myelin proteolipid protein in cadavers with multiple sclerosis (Lee et al., 2014). It has also been used to identify in vivo myelin degeneration in patients with schizophrenia (Ganzetti et al., 2015; Iwatani et al., 2015), multiple sclerosis (Beer et al., 2016), and bipolar disorder (Ishida et al., 2017). Fur–ther, the method has also been used to demonstrate that higher estimated myelin within the cerebral cortex is cor–related with performance stability on speeded tasks (Grydeland et al., 2013).

Although we are not aware of any research investigat–ing the association between sub-cortical myelin content, as measured by T1w/T2w, and cognitive performance, it would be expected that lower myelination white matter tracts would be associated with lower PS in cognitively healthy individuals. Moreover, since age-related decrease in myelination has been clearly demonstrated (Bartzokis, 2004), it would be predicted that older individuals would present with lower myelination levels than younger indi–viduals and that this difference in myelination would be associated with a slower PS. Thus, the aim of this study was to investigate whether the myelin content of major white matter tracts, assessed with the T1w/T2w ratio, was predictive of PS in a large sample of cognitively healthy middle-age and older adults.

## 2. Materials and Method

### 2.1. Participants

Participants were selected from the MRI sub-study within the PATH Through Life Project (PATH) which has been described in detail elsewhere (Anstey et al., 2012). Briefly, PATH is an ongoing population-based longitudi–nal study that aims to track the course of cognitive abil–ity, mental health disorders, substance use and dementia across the lifespan. Participants are randomly selected from the electoral roll of the Australian Capital Territory and surrounding regions. Data collection started in 1999 and participants are reassessed every four years.

The PATH study consists of three cohorts: 2024 years (young adult), 4044 years (middle-age), and 6064 (older-age) years at baseline. The focus of this study is on the middle-age (MA) and older-age (OA) cohorts at the third assessment, due to the availability of higher quality T1-w and T2-w MRI scans for both the MA and OA partici–pants at this time-point. Of the 2530 MA and 2550 OA participants recruited into the study, 304 MA and 303 OA participants had complete imaging data at the third assessment. However, 14 scans were excluded due to poor qual–ity. From this sample, a further 23 participants were ex–cluded due to: epilepsy (n=2), having a history of stroke (n=14), Parkinsons disease (n=3), dementia (n=2) and cognitive impairment (n=2) as defined by a Mini-Mental Status Exam score of less than 25. The final sample avail–able for analysis included 570 participants (277 MA and 293 OA). The selected sample did not differ significantly from the overall MA and OA PATH cohort on sex and education; however, it was significantly younger (p = .048).

### 2.2. Socio-demographic, health and genetic measures

Years of education, alcohol consumption, smoking, physical activity were assessed using self-report. Alco–hol consumption was assessed as the number of standard alcoholic drinks consumed per week (Alcohol Use Dis–orders Identification Test; Babor et al., 2001). Physical activity was assessed as the number of hours per week of mild, moderate and vigorous exercise. To provide an intensity-sensitive continuous score of physical exer–cise, the three levels of activity were combined using a weighted procedure such that hours of mild physical ac–tivity were multiplied by 1, hours of moderate physical activity by 2 and hours of vigorous physical activity by 3(Lamont et al., 2014). Depressive symptomology was assessed using the Goldberg Depression Score (Goldberg et al., 1988). Seated systolic and diastolic blood pres–sures (BP) were averaged over two measurements after a 5-minute rest and participants were classified as hyperten–sive if they were on medical therapy for hypertension or if they had an average systolic BP 140mm Hg or diastolic BP 90mm Hg. Genomic DNA was extracted using cheek swabs and was used to identify the presence of APO*e4 genotype (Christensen et al., 2008).

### 2.3. Measures of cognitive PS

PS was assessed using five different tasks. The Sym–bol Digit Modalities Test (SDMT; Strauss et al., 2006), was scored as the number of correct matches identified according to the stimulus symbol digit code, within a 90–s period. Simple (SRT) and choice reaction time (CRT) were assessed by giving participants a small box to hold with both hands, with left and right buttons at the top to be depressed by the index fingers. The front of the box had three lights: a red stimulus light under each of the left and right buttons, and a green get-ready light in the middle. For SRT task, participants placed their right hand, on the right button and were asked to press it as quick as possi–ble when they saw the red stimulus light up. For the CRT task, participants were asked to place their right finger on the right button and their left finger on the left button and to press the corresponding button when the left or right red light lit up. There were 4 blocks of 20 trials measur–ing SRT, followed by two blocks of 20 trials measuring CRT. The mean reaction time was the average across all trials. Trial Making Task Part A (TMT-A; Reitan, 1958) was scored as the amount of time taken to complete the task and the Purdue Pegboard task using both hands (PP; Tiffin and Asher, 1948) was scored as the number of pairs of pins placed into the pegboard device within 30-s.

### 2.4. MRI data acquisition

All participants were imaged in a 1.5-Tesla Siemens Avanto scanner (Siemens Medical Solutions, Erlangen,Germany). T1-w images were acquired in sagittal orienta–tion with (repetition time/echo time/flip angle/slice thick–ness = 1160ms/4.17ms/15/1 mm) matrix size 256 256 and voxel size of 1 × 1 mm. T2-w images were acquired in coronal orientation with (repetition time/echo time/flip angle/slice thickness = 9680ms/115ms/150/4 mm) matrix size 256 × 256 and voxel size 0.898 × 0.898 mm.

### 2.5. MRI data analysis

T1-w and T2-w image were pre-processed and com–bined, following the method and workflow outlined in Ganzetti et al. (2014, 2015). This process included bias correction and intensity calibration on both the T1-w and T2-w image before they were combined. This entire process was undertaken us–ing Freesurfer, FSL and the MINC imaging tool-box(http://www.bic.mni.mcgill.ca/ServicesSoftware).

First the T1-w images were transformed using the MNI152 atlas into Talairach space. A rigid-body trans–form was then used to match the T2-w image to the al–ready transformed T1-w image. To address intensity bias due to distortions in the B1 field between T1-w and T2-w images, each image was first individually bias-corrected using the mri_nu_correct tool from the MINC imaging toolbox (http://www.bic.mni.mcgill.ca/ServicesSoftware) with the default setting. As the T1w/T2w technique is a qualitative technique, it is susceptible to intensity scaling discrepancies across both individuals and scanners. As such, a calibration procedure recommended by Ganzetti et al. (2014) that involved the linear transformation of the bias-corrected images was implemented. Specifically, two non-brain areas of homogenous intensity were se–lected: one area that contains relatively high values in the T1-w scan and relatively low values in the T2-w scan, and another area with the reverse characteristics. Con–sistent with Ganzetti et al. (2014), the temporal muscle and the eyeball humour were selected. To calculate the scaling factors, the mode value in each of the selected ar–eas was extracted and compared with corresponding val–ues from the high-resolution International Consortium for Brain Mapping (ICBM) reference image. The T1-w and T2-w images were then separately multiplied by the re–sulting scaling factor to create the calibrated images. Af–ter calibration, the T1-w image was divided by the T2-w image to create the final T1w/T2w ratio image.

### 2.6. Regions of interest

Consistent with previous research (Ganzetti et al., 2014), a total of 6 white matter tracts with putatively high myelin content were selected: anterior corona radiata (ACR), superior corona radiata (SCR), pon–tine crossing tract (PCT), anterior limb of the internal capsule (ALIC), genu of the corpus callosum (GCC), splenium of the corpus callosum (SCC). In addition, three areas putatively low in myelination (caudate, putamen and thalamus) were selected as compassion re–gions to confirm that the T1w/T2w measure was cor–rectly discriminating between high and low myelinated regions. All ROIs were defined using the stereotaxic John Hopkins University white-matter tractography atlas (http://cmrm.med.jhmi.edu; Oishi et al., 2008).

### 2.7. Statistical analysis and experimental design

Statistical analyses were computed using IBM SPSS Statistics 24. Age was split into two variables to sepa–rately assess the within and between cohort variability in age: age group (AgeG) and age centred (AgeC). AgeG in–dicated whether the participant belonged to the MA or OA group. AgeC was calculated by subtracting the rounded age of the youngest participant from the exact age of par–ticipants in each group. After being converted to z-scores, the five independent tests of PS were subject to a princi–pal components analysis (PCA) in order to extract a sin–gle common factor of PS. Principal components with an eigenvalue greater than 1 were retained for further con–sideration. The selected principal component of PS was confirmed by the high and consistent loadings of individ–ual measures of PS contributing to it.

Independent sample t-tests and chi-squared tests were used to assess AgeG differences between the MA and OA groups on socio-demographic, health, genetic variables. Multiple paired sample t-tests were used to assess whether each of the selected white matter ROIs had higher mean T1w/T2w values from each of the grey matter ROIs. Mul–tiple ANCOVAs were used to identify AgeG and Sex dif–ferences and interactions in myelin content, controlling for AgeC, within the six white matter tracts. The same approach was used to identify AgeG and Sex differences in the principal component of PS.

To assess whether tract myelination could predict PS, hierarchical linear regression analyses were performed to examine the association between myelin content, and PS. The preliminary model included AgeG, AgeC, Sex and Education as independent variables and PS as the de–pendent variable (block 1). Myelin content of the ROI and a lateralization index [calculated with the formula [(L-R)/(L+R)*100] were entered as an independent vari–able (block 2). To determine whether mean ROI inten–sity could account for any additional variance in PS over health and genetic variables, block 3 included all variables examined. The lateralization index was only retained in the models if it was a significant predictor. All two-way and three-way interactions terms were examined (block 4). Significance was set at p ¡ 0.05 and correction for mul–tiple comparisons was applied using the Holm-Bonferroni method (Holm, 1979).

## 3. Results

### 3.1. Sample characteristics

Group comparison between the OA and MA groups re–vealed that the OA group had fewer females, was significantly less educated, had a higher physical activity score, higher rates of hypertension and scored higher on depres–sive symptomology (Table 1).

**Table 1.**
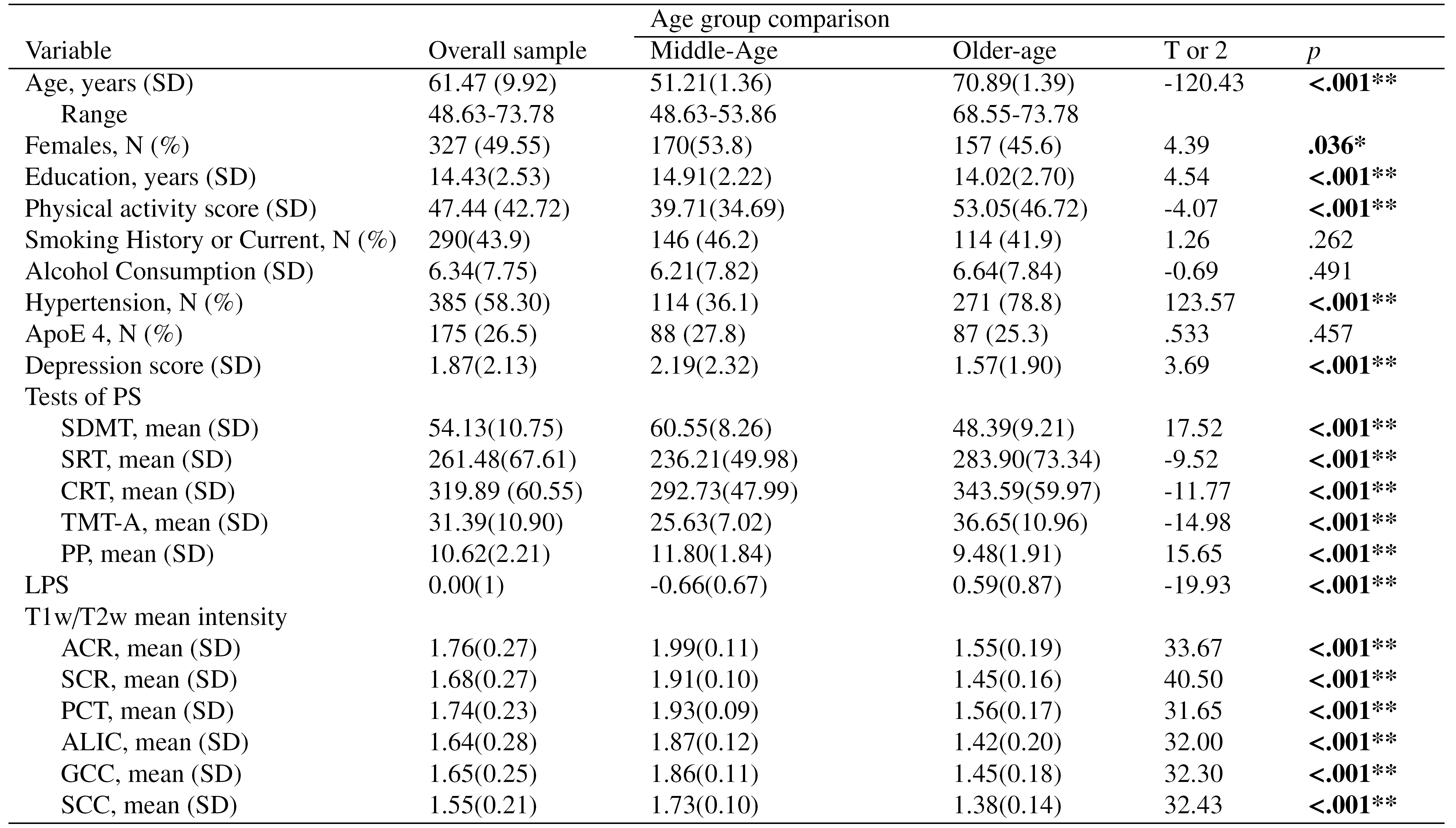
Sample characteristics and age-group differences.

### 3.2. Comparison ofT1w/T2w method across high and low myelinated ROIs

As predicted, selected white mater tracts had signifi–cantly higher mean T1w/T2w values compared to the grey matter ROIs (p < .001), with a mean difference rang–ing from .174 (11.25%) to .577 (32.75%) and SD rang–ing from .101 to .239 with large effect sizes (Cohens d = .733 to 2.12). Mean T1w/T2w values across all ROIs are shown in Figure 1.

**Figure 1.**
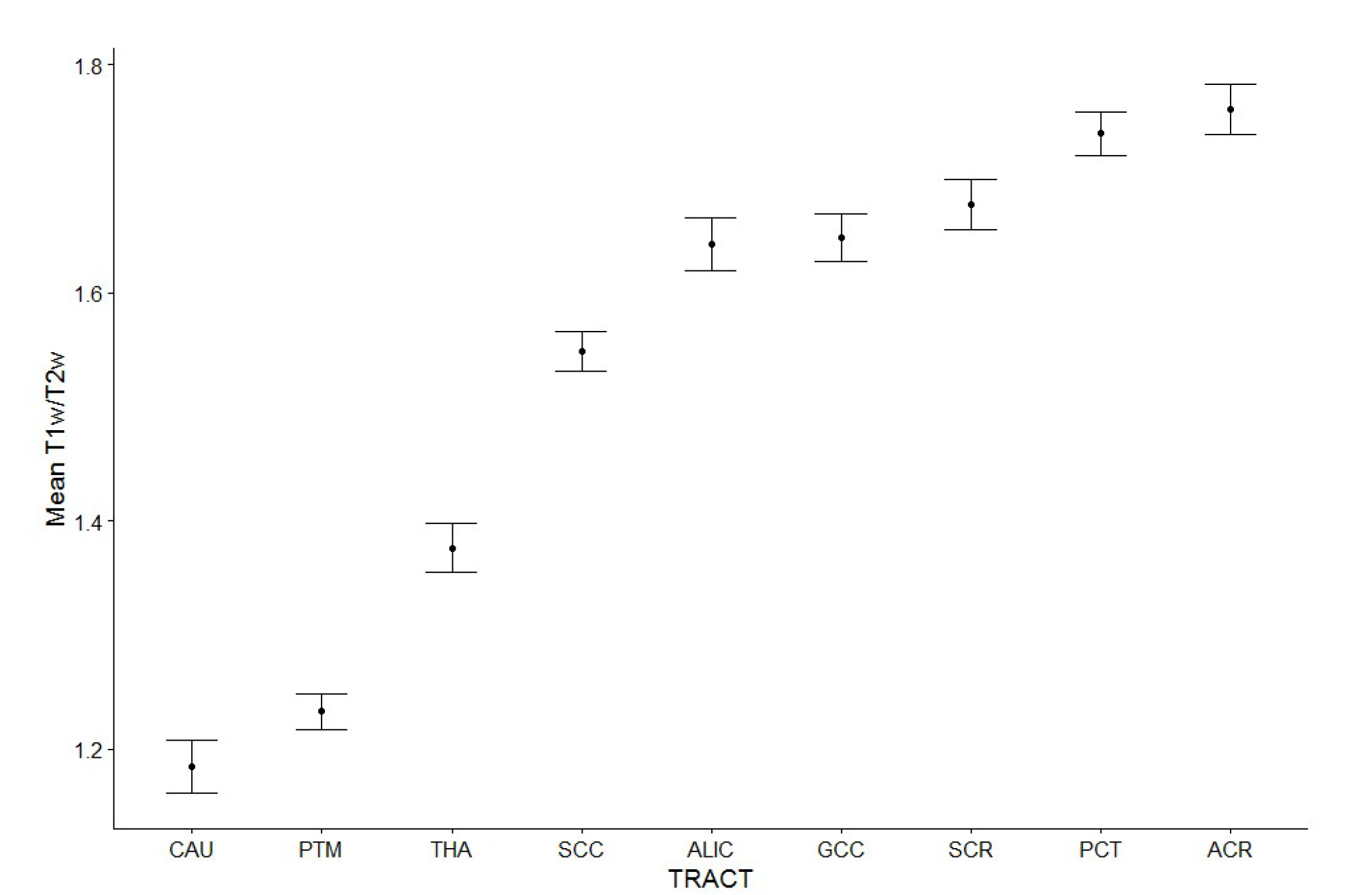
Mean and confidence interval of T1w/T2w values for the three grey matter and six white matter ROIs. As expected grey matter areas had lower mean values than white matter areas. Abbreviations: ACR = anterior corona radiata; SCR = superior corona radiata; PCT = pontine crossing tract; ALIC = anterior limb of the internal capsule; GCC = genu of the corpus callosum; SCC = splenium of the corpus callosum.

### 3.3. Age group and sex differences in tract myelination

The Tract by AgeG by Sex ANCOVA revealed the fol–lowing effects. A main effect for AgeG was detected, in–dicating that the OA group had significantly lower myelin content in all six of the white matter tracts examined (Table 1), with large effect sizes: 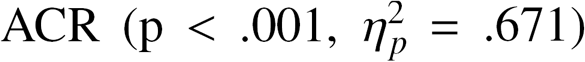, 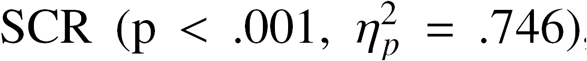, PCT 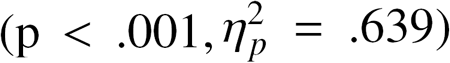, 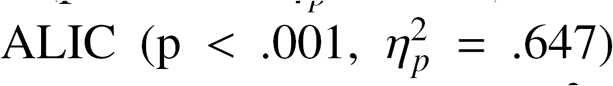, 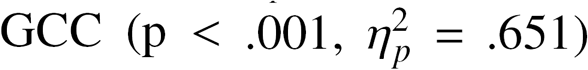 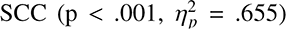. On average, the white matter tracts of the OA group were 21.95% less myelinated when compared to the MA group, with the biggest difference seen in the SCR (24.08%) and the ALIC (24.06%), followed by ACR (22.11%), GCC (22.04%) SCC (20.23%) and PCT (19.17%).

### 3.4. Age as a predictor of PS

A significant main effect of sex was found within the 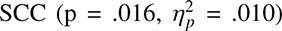, with females being 1.60% more myelinated than males. An AgeG by sex interaction was detected in the 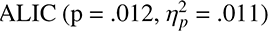, 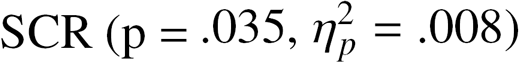, 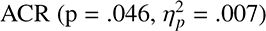 and the PCT 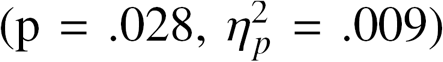. In all cases, females had higher levels of myelination in the MA group, but lower in the OA group. Figure 2 shows age and sex differences in tract myelination.

**Figure 2.**
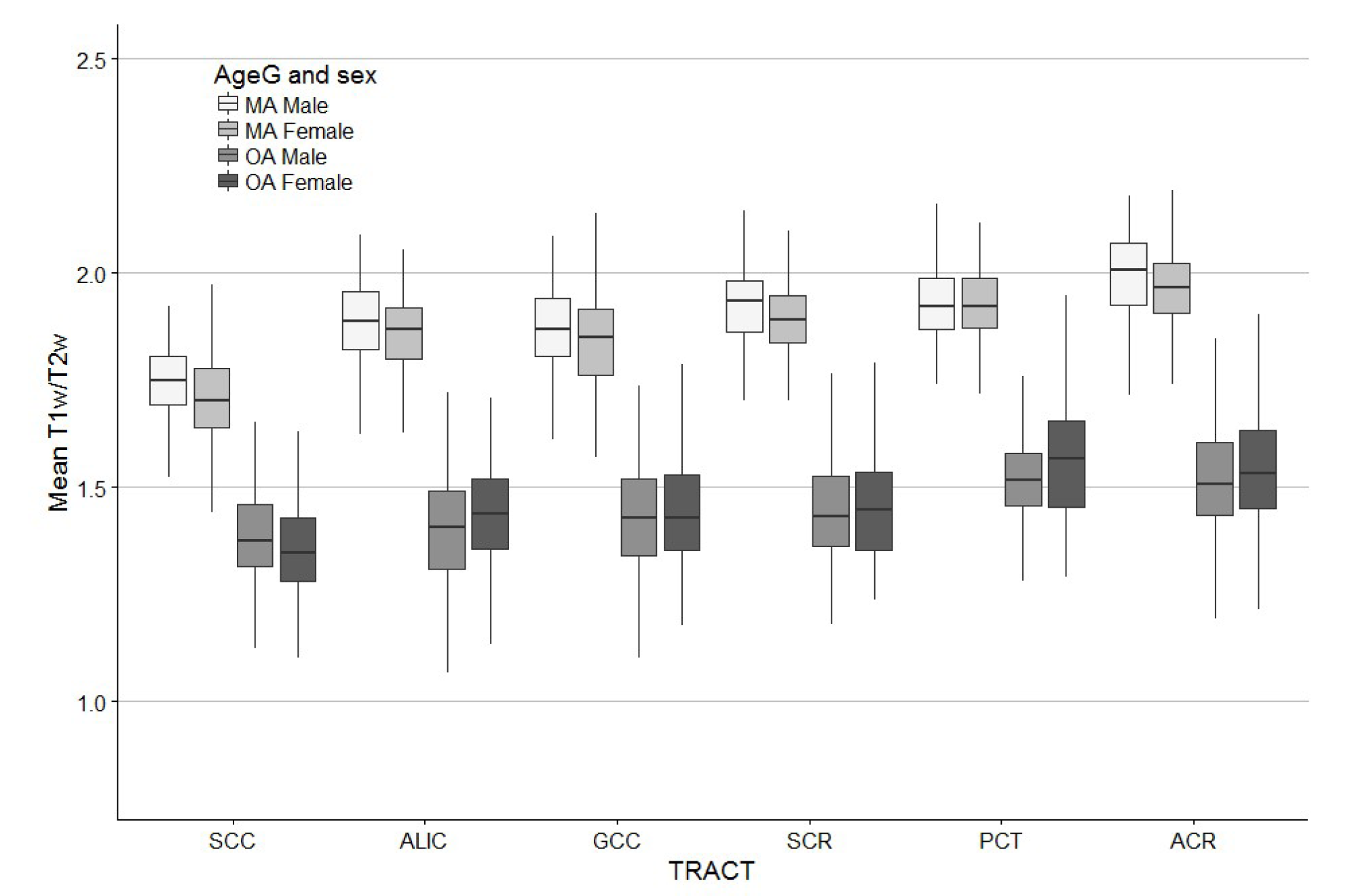
Age group (AgeG) and sex differences in estimated tract myelination. The boxplot shows that on average middle-ages partic–ipants (MA) had higher estimated myelination levels than older-aged participants (OA) and that while females had higher estimated myeli-nation levels in MA within ALIC, SCR and ACR, they presented with lower levels in OA within these three tracts. ACR = anterior corona radiata; SCR = superior corona radiata; PCT = pontine crossing tract; ALIC = anterior limb of the internal capsule; GCC = genu of the corpus callosum; SCC = splenium of the corpus callosum.

### 3.5. Principal component of PS

A principal component analysis (PCA) was run on the five PS tasks in order to extract a single component of PS. The PCA revealed one factor that had an eigenvalue greater than one ( = 2.86) and which explained 57.50% of the total variance. Loadings for each test were as fol–lows (with communalities in parentheses): CRT = .842 (70.9%), SDMT = -.803 (64.4%), SRT = .761 (57.9%), TMT-A= .721 (52.0%) and PP = -.650 (42.3%). Con–sequently, as expected this component was interpreted as reflecting a latent factor of PS (LPS).

### 3.6. Age group and sex differences in PS

The AgeG by Sex ANCOVAs testing performance on LPS revealed a significant main effect of AgeG, with the OA group performing slower on average, with a large ef–fect size 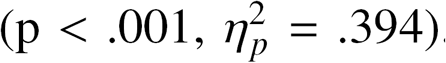. No significant main ef–fect for Sex or interaction effects were detected. To deter–mine whether individual measures contributed differently to these effects, follow-up analyses revealed a significant main effect of AgeG on all individual tests of PS: SDMT 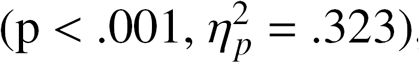, Trails 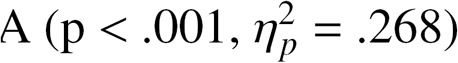, SRT 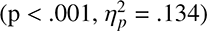, 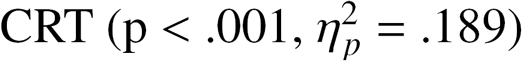, PP 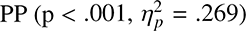. Significant main effect of Sex revealed that females performed faster for 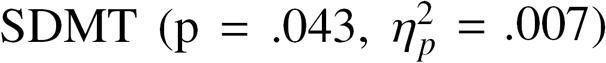 and Pegboard 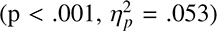, and slower for 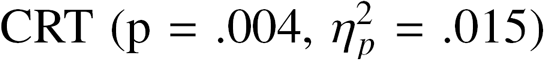 and 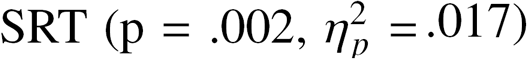. The differences in Pegboard, CRT and SRT sur–vived correction for multiple comparisons. No significant interaction effects were detected.

Hierarchical regression modeling revealed that in–creased myelin content significantly predicted faster PS but only in the ALIC (Table 2). One SD increase in the myelin content of the ALIC was associated with 2.53% increase in PS. This was a robust finding as the associa–tion remained significant after controlling for all covariates and after correction for multiple comparisons. Associations in all other white matter tracts followed a similar trend but did not reach significance. No significant inter–actions were detected. Scatterplots of PS as a function of myelin content for each tract are presented in Figure 3.

**Table 2:**
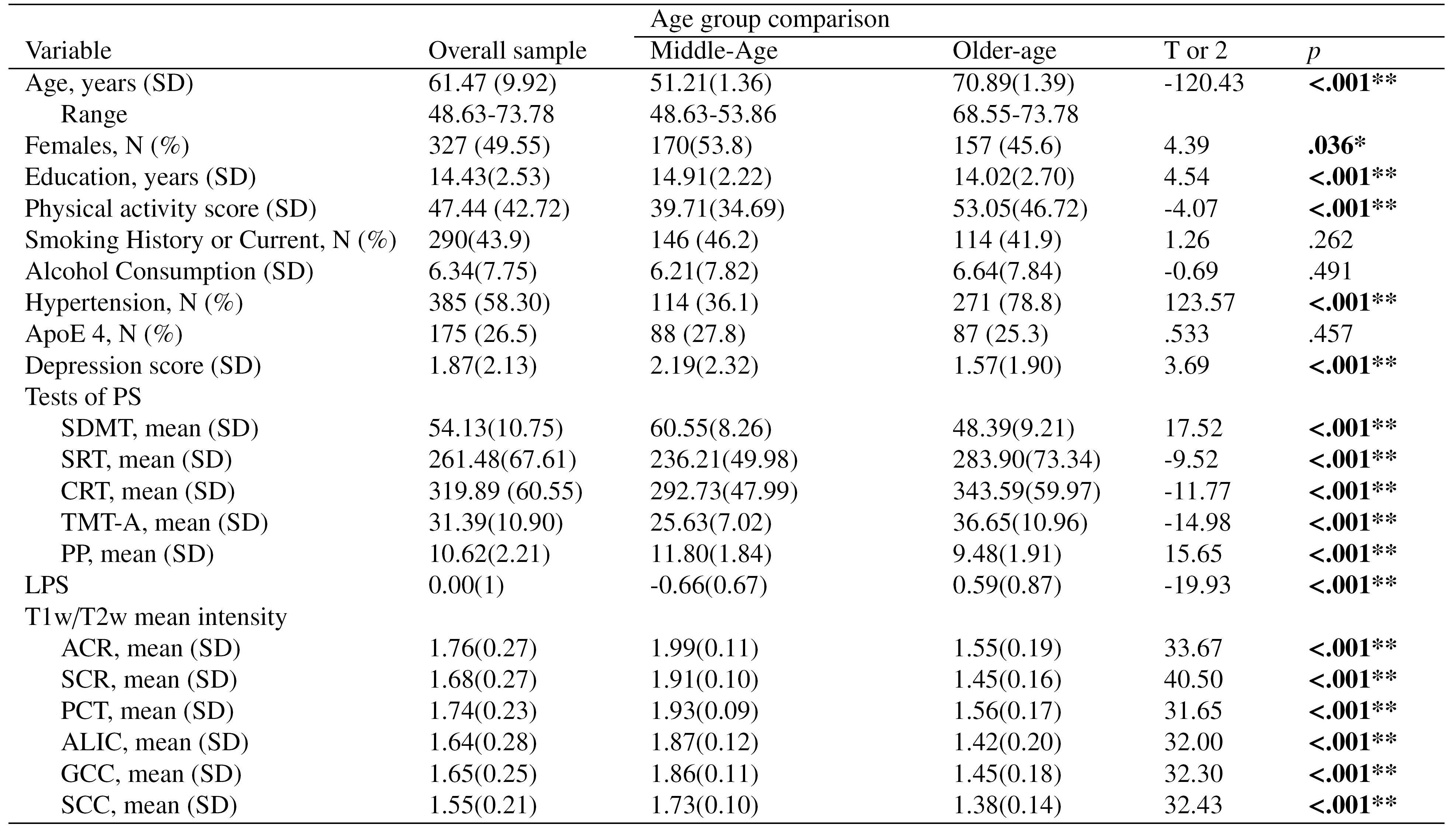
Abbreviations: PS = processing speed; LPS = latent factor of processing speed; SDMT = Symbol Digits Modalities Test; SRT = simple reaction time; CRT = choice reaction time; TMT-A = Trial Making Task A, PP = Purdue Pegboard; ACR = anterior corona radiata; SCR = superior corona radiata; PCT = pontine crossing tract; ALIC = anterior limb of the internal capsule; GCC = genu of the corpus callosum; SCC = splenium of the corpus callosum.* Significant at p < .05, *** Significant at p < .001

**Figure 3.**
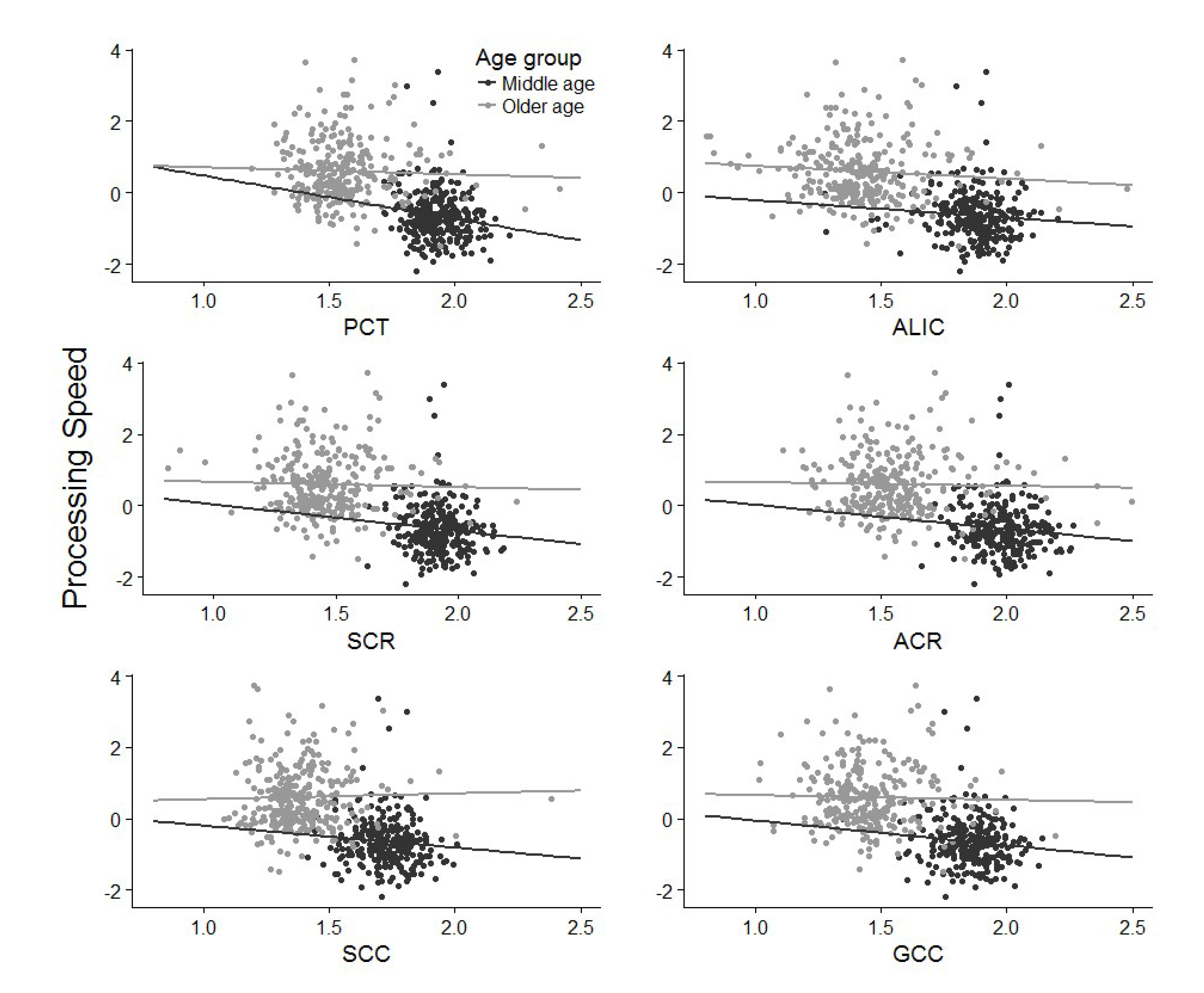
Scatterplots of processing speed component as a function of estimated myelin content within each of the six selected tracts. ACR = anterior corona radiata; SCR = superior corona radiata; PCT = pontine crossing tract; ALIC = anterior limb of the internal capsule; GCC = genu of the corpus callosum; SCC = splenium of the corpus callosum.

### 3.7. Age as a predictor of PS

AgeG remained a significant predictor of PS in all six of the final models, with membership of the OA group re–sulting in a 16.56% to 19.42% slower PS depending on which tract was entered. The effect of AgeG survived correction for multiple comparison in all models. AgeC also remained a significant predictor in all final models, indicating that each one year increase in age within age groups was associated with a 0.95% to 1.03% decrease in PS, depending on which tract was entered into the models. However, this effect did not survive correction for mul–tiple comparisons. No significant interactions between AgeG, AgeC were detected.

### 3.8. Sex as a predictor of PS

Sex was not a significant predictor in any of the six models. However, a trend suggested females performed 1.23% to 1.59% faster than males, depending on which tract was entered into the models. No significant interac–tions involving sex were detected.

#### 3.9. Post-hoc analyses

##### 3.9.1. Hemispheric differences in myelin content

In order to better understand the role of myelination in PS, we conducted further analyses to investigate whether there were hemispheric differences in myelin content and whether hemispheric asymmetries contribute to the effects detected above. Analyses revealed a significant main ef–fect for hemisphere within all tracts with the left hemi–sphere being less myelinated than the right within the 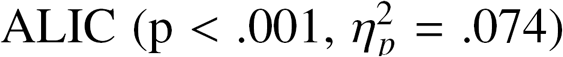, 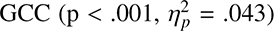 and 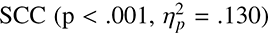, whereas the reverse was found within the 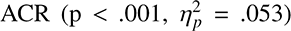, 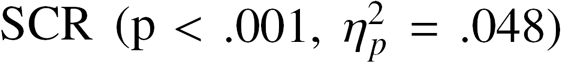 and 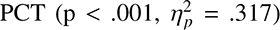. A significant hemisphere by AgeG interaction effect found within the 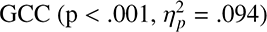 revealed that while the right hemisphere was less myelinated within the MA group, the left hemisphere was less myelinated within the OA group. A significant hemisphere by sex interaction was also found within the 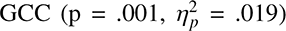, with the left hemisphere being more myelinated within fe–males, whereas no significant difference was found within males. All significant main effects and interactions for hemisphere survived correction for multiple comparisons.

##### 3.9.2. Hemispheric asymmetries in myelin content as a predictor of PS

The laterality index of the SCC (β = -.018, p = .002) was a significant predictor of PS. As such, left and right SCC were examined in separate regression models (Table 3). Analyses revealed that the left SCC was a significant predictor of PS. One SD increase in the myelin content of the left SCC was associated with 2.20% increase in PS. This was a robust finding as the association remained sig–nificant after controlling for all covariates and survived correction for multiple comparisons. The right SCC was not a significant predictor in either models, and no signif–icant interaction effects were detected.

**Table 3.** ^a^ Model includes AgeG, AgeC, Sex and Education ^b^ Model includes myelin content (mean T1w/T2w) of ROI and the laterality index. As the laterality index was dropped from the model unless it was a significant predictor, it was removed from all models except for SCC models ^c^ Model includes alcohol consumption, smoking, physical activity, hypertension, presence of APO*E4 genotype, depressive symptomology ** Significant at p < .01

| ROI | Model 1<sup>a</sup> | Model 2<sup>a,b</sup> |  |  | Model 3<sup>a,b,c</sup> |  |  |
| --- | --- | --- | --- | --- | --- | --- | --- |
| | $R^2$ | $\Delta R^2$ | $\beta$ | p | $\Delta R^2$ | $\beta$ | p |
| ACR | .421 | .002 | -.340 | .158 | .014 | -.362 | .132 |
| SCR | .410 | .002 | -.353 | .148 | .014 | -.363 | .137 |
| PCT | .411 | .004 | -.394 | .063 | .014 | -.405 | .052 |
| ALIC | .410 | .007 | -.519 | **.009\*\*** | .014 | -.526 | **.008\*\*** |
| GCC | .410 | .003 | -.387 | .081 | .014 | -.433 | .050 |
| SCC | .418 | .011 | -.275 | .319 | .015 | -.304 | .270 |

##### 3.9.3. Partial correlations between myelin content and individual tests of PS

To determine which individual tests of PS were best predicted by the myelin content of the ALIC and left SCC, post-hoc partial correlations the between the two tracts and the five different tests of PS were computed, control–ling for AgeG, AgeC, Sex and Education. To assump–tions of normality and assess relative contributions, the z-scores for each test were used. Partial correlations re–vealed that, SDMT was the test most strongly correlated with the myelin content of the ALIC (r = .093, p = .037), followed by CRT (r = -.073, p = .103), SRT (r = -.050, p = .264), TMT-A (r = -.054, p = .231) and PP (r = -.008, p = .852). Whereas for the left SCC, TMT-A was most strongly correlated (r = -.121, p = .007), followed by CRT (r = -.108, p =.016), SRT (r = -.089, p = .047), PP (r = .057, p = .203) and SDMT (r = .049, p = .273).

## 4. Discussion

### 4.1. Myelin content as a predictor of PS

The present study investigated whether myelin content estimated in vivo within major white matter tracts is pre–dictive of PS in a sample of healthy community-dwelling adults and whether lower myelin content could be de–tected in older compared to younger individuals. The main finding was that higher myelin content in the ALIC and the left SCC significantly predicted faster PS scores and that myelin content was lower in older individuals.

**Table 4:**
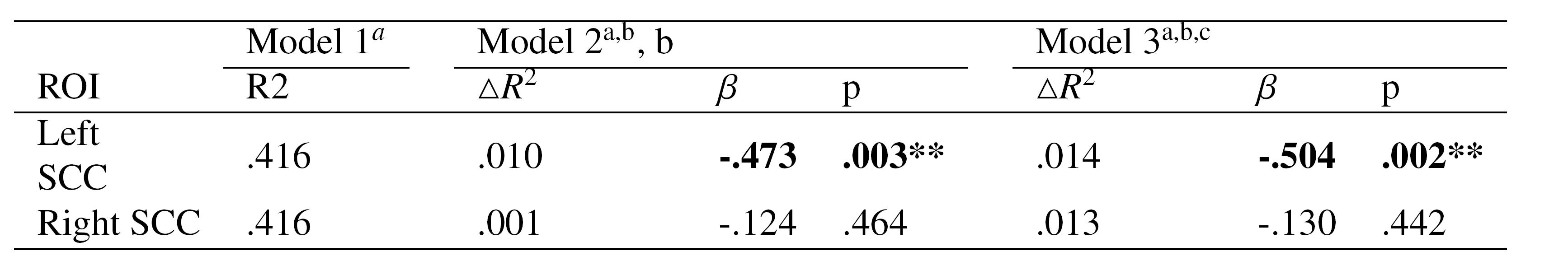
^a^ Model includes AgeG, AgeC, Sex and Education ^b^ Model includes myelin content (mean T1w/T2w) of ROI ^c^ Model includes alcohol consumption, smoking, physical activity, hypertension, presence of APO*E4 genotype, depressive symptomology ** Significant at p < .01

| ROI | Model 1<sup>a</sup> | Model 2<sup>a,b</sup>, b |  |  | Model 3<sup>a,b,c</sup> |  |  |
| --- | --- | --- | --- | --- | --- | --- | --- |
| | R<sup>2</sup> | $\Delta R^2$ | $\beta$ | p | $\Delta R^2$ | $\beta$ | p |
| Left SCC | .416 | .010 | **-.473** | **.003\*\*** | .014 | **-.504** | **.002\*\*** |
| Right SCC | .416 | .001 | -.124 | .464 | .013 | -.130 | .442 |

### 4.1.1. T1w/T2w as an index of myelin content

Our analyses are in agreement with previous research (Ganzetti et al., 2014, 2015), confirming that myelin con–tent estimated with the T1w/T2w ratio is significantly lower in grey matter ROIs compared to white matter ROIs, where myelin is evidently found in higher concentrations in histological studies. Specifically, our findings are con–sistent in showing that within the grey matter ROIs, the thalamus was the most myelinated structure (Whittall et al., 1997; Mdler et al., 2008; Ganzetti et al., 2015). Simi–larly, our findings within the corpus callosum are consis–tent with previous research and histology studies in show–ing that the genu has higher myelin content than the splenium (Aboitiz et al., 1992; Lee et al., 2014). However, while past studies have found the ALIC to have the high–est myelin content within white matter areas (Whittall et al., 1997; Ganzetti et al., 2014), we found the ACR to have the highest estimated myelin content. As the sample used in the current study is substantially narrower and older than that of previous studies, it is possible this discrep–ancy represents an ageing effect. Supporting this hypoth–esis, we found that the ALIC showed one of the largest estimated difference in myelination (24.06%) between the OA and MA group. In addition, the MA group had signif–icantly higher T1w/T2w values in all six tracts, which is consistent with the previously reported trajectory of brain myelin content which begins to decline soon after mid–life (age 35-40; Bartzokis et al., 2003; Bartzokis et al., 2010). It should be noted that while previous literature and the above findings suggests that T1w/T2w signal is largely dependent on myelin content, it cannot be ruled out that axonal fibre loss contributes to this signal.

### 4.1.2. Myelin content predicts PS

Lower myelination estimated with the T1w/T2w ratio is present in clinical groups, and higher estimated myelination within the cortex is correlated with performance stability on speeded tasks (Grydeland et al., 2013). To our knowledge this is the first study to utilise the T1-w/T2-w ratio to demonstrate an association between sub-cortical myelin content and PS in generally healthy community-living individuals. These results support the claim that the increased speed of signal transmission provided by myelin can predict better outcomes within the cognitive domain of PS, even in healthy adults who are not likely to have marked myelin degradation similar to those found in clinical samples. Further, the results found were ro–bust as they remained significant even after controlling for socio-demographic, health, genetic covariates and multi–ple comparisons.

The association between higher estimated myelin con–tent and faster PS was found to be significant in the ALIC and left SCC and consistent trends were found in all tracts investigated. The ALIC is known for high myelin content, connecting the prefrontal cortex to thalamic nuclei, and the motor cortex to the anterior horn of the spinal cord (Mai et al., 2015). While the cognitive tests used within the current study primarily measure PS, they do not assess this property in relation to a single function and may re–flect axonal conduction contributing to a variety of motor, perceptual and cognitive processes. As such, demyelination of the ALIC may result in the disruption of cogni–tive processes reliant on different circuits. Thus, since the cortico-thalamic circuit contributes to a range of cognitive processes that include learning and memory, inhibitory control, decision-making, and the control of visual orient–ing responses (Haber and Calzavara, 2009), differences in myelination of its fibres may modulate performance of these processes. Alternatively, since the cortico-spinal circuit forms the major motor control pathway, the slow–ing of signals travelling through this circuit could result in the slowing of PS and psychomotor response seen in the current study. In support of this, demyelination and ax–onal damage within the ALIC has been linked with motor impairment in multiple sclerosis patients (Lee et al., 2000) and PS deficits in non-demented older adults (O’brien et al., 2002). Importantly, the speed at which an electrical signal travels along an axon is directly related to the de–gree and quality of myelination (Seidl, 2014). Tempo–ral efficacy is essential within neural circuits to ensure computations are completed on time and synchronously. Evidence suggests that myelin from later-differentiating oligodendrocytes, such as that found in the ALIC, is less effective and more vulnerable to the age-related effects of inflammation and oxidative stress (Brickman et al., 2012; Kohama et al., 2012). If neural efficacy is compromised due to degradation of myelin sheath in vulnerable areas, like the ALIC, it may result in cognitive and behavioural slowing, such as that seen in the current study.

In addition to the ALIC, the myelin content of the left SCC significantly predicted PS. The SCC is a major com–missural tract, accommodating interhemispheric connec–tions between visual, parietal and posterior cingulate ar–eas (Knyazeva, 2013), and is closely associated with PS, in that age-related volume loss (Anstey et al., 2007) and white-matter hyper-intensities within the SCC have been shown to be associated with slower speed of processing (Park et al., 2014). These age-related findings may be due to increased interhemispheric synchronisation facilitated by the heavily myelinated fibres of the SCC (Hinkley et al., 2012). As such, the current study suggests that age-related PS disruptions associated with the SCC may be due to myelin degradation within the structure. However, only the myelin content estimate of the left SCC was a significant predictor of PS. While it is possible that this lateralised finding is due to noise or measurement error, the left hemisphere, and in particular the left SCC may be more prone to neurodegeneration (Yoon et al., 2011). Ad–ditionally, in Alzheimers disease, cortical atrophy begins earlier and progresses faster within the left hemisphere (Thompson et al., 2007). Consequently, the laterality ef–fect observed in the present study may reflect a greater vulnerability of the left SCC to the adverse effects of ageing.

It is of clinical utility to determine which of the five in–dividual tests of PS would be best predicted by myelin deficits in the ALIC and left SCC. Post-hoc analysis showed that the SDMT and TMT-A were most strongly correlated with the estimated myelin content of the ALIC and left SCC respectively. One of the major aspects that differentiates these two tests from the others is increased cognitive complexity as opposed to SRT, CRT, and PP which are primarily reliant on precise motor skills and vi–sual feedback. This is consistent with the fact that the ALIC contributes to the fronto-thalamic circuitry which is involved in complex cognition such as executive func–tions.

### 4.1.3 Strengths and limitations

One of the primary strengths of this study was the ro–bust measurement of PS. By extracting a latent measure of PS from five different individual tests, unwanted variance relating to other properties was minimised by analysing common aspects of tasks that vary in methodology but load heavily on PS (Cepeda et al., 2013). In addition, the large sample size and carefully selected covariates were also substantial strengths.

Despite this, the narrow age-range of the two groups and the relative good health of our sample may have re–stricted variance in myelination that might be otherwise expected in a population with a greater age range or in a clinical sample. Additionally, due to the cross-sectional design used, causal inferences on the association between T1w/T2w signal and PS cannot be drawn. Future stud–ies should therefore apply this technique to longitudinal datasets to determine both lifelong myelination trajecto–ries and test their associations across a greater range of cognitive domains such as memory and attention which may also be materially affected by progressive age-related demyelination.

It should also be noted that while the measure of myelination used in this study has shown some concor–dance with immunostaining for myelin proteolipid protein (Nakamura et al., 2016), it is yet to be directly validated against histological measures. However, a recent study demonstrated that within a small sample, the measure had high test-retest reliability (Arshad et al., 2017). Future studies should continue to aim to directly validate the ori–gin of the T1w/T2w signal by using histology techniques in animal and cadaveric models.

### 4.1.4 Conclusion

Estimating brain myelination using the T1w/T2w tech–nique is relatively easy to implement and does not re–quire lengthy acquisition times or a complex processing pipeline; therefore, it is a practical method to identify age-related brain changes and should be further investigated in future research as a biomarker for neurocognitive dis–eases. Using this technique, the current study is the first to demonstrate *in vivo* that higher estimated myelin content of white matter tracts assessed with a specific and sensi–tive measure is associated with faster PS in healthy adults, even after controlling for socio-demographic, health and genetic variables.

